# Replication stress tolerance and management differs between naïve and primed pluripotent cells

**DOI:** 10.1101/2022.05.12.491744

**Authors:** Georgia R. Kafer, Aisling O’Connor, Samuel Rogers, Pierre Osteil, Christopher B. Nelson, Hilda A. Pickett, Patrick P.L. Tam, Anthony J. Cesare

## Abstract

Replication stress is an endemic threat to genome stability. For reasons unknown, replication stress response factors become essential during peri-implantation development. This coincides with a stem cell potency switch from the naïve to the primed state. Using genetically matched, chimera-derived mouse naïve embryonic (mESC) and primed epiblast stem cells (mEpiSC) we found that replication stress management differs between potency states. Primed mEpiSCs rely on Atr activity to prevent replication catastrophe, minimize genomic damage, avoid apoptosis, and re-enter the cell cycle. Conversely, under replications stress, mESCs readily activate Atm regardless of Atr activity, undergo replication catastrophe, and induce apoptosis. Primed pluripotent cells therefore engage Atr to counteract replication difficulties and maintain viability, whereas cells in the naïve state are more readily cleared under the same conditions. We anticipate these divergent strategies enable pluripotent cells of different potency states to meet associated proliferative or developmental demands during early development.

## Introduction

Maintaining genome integrity during embryogenesis is critical for fetal survival and long-term health. A unique challenge of early development is protecting the genome while maintaining rapid cell proliferation. Embryonic pluripotent cells progress through a series of programmed potency changes including the naïve state of mESCs within the blastocyst Inner Cell Mass (ICM), and the primed state of mEpiSCs within the gastrulating embryo (Nichols and Smith, 2009; Weinberger et al., 2016). The potency switch from naïve to primed cells corresponds temporally with a pre-gastrulation phase referred to as peri-implantation development (Rossant and Tam, 2009). Average proliferation rates in naïve and primed pluripotent cells are 12 and 4 hours, respectively; a pace substantially faster than the typical 24-hour doubling times observed in rapidly dividing somatic and cancer cells (Bolton et al., 2016; Singla et al., 2020; Snow, 1977; Starostik et al., 2020). While rapid cell proliferation correlates with mutation and genomic instability in somatic and oncogenic tissues, pluripotent embryonic cells *in vivo,* and in culture, maintain expeditious cell cycles and a 1000-fold lower mutational burden in comparison to differentiated tissues (Giachino et al., 2013; Hong et al., 2007; Tichy, 2011; Xiong et al., 2015). Pluripotent cells must therefore employ robust cellular strategies to maintain overall genome health of the conceptus whilst also facilitating rapid proliferation.

Compromised DNA replication, termed replication stress, is the primary genomic insult associated with rapid cell division cycles (Zeman and Cimprich, 2014). Replication stress in somatic cells promotes excessive single stranded DNA (ssDNA) at replication forks. Replication Protein A (Rpa) binds this ssDNA, promoting localized recruitment of the master replication stress response regulator, ataxia telangiectasia and Rad3-related (Atr) kinase (Zou and Elledge, 2003). Following double strand break (DSB) induction, the related ataxia telangiectasia mutated (Atm) kinase localizes to DNA lesions and regulates the DSB response (Paull, 2015). Somatic Atr inhibition coupled with replication stress can exhaust the cellular Rpa pool, exposing ssDNA at stalled forks to breakage, thereby acutely activating Atm (Toledo et al., 2017; Toledo et al., 2013). This event cascade, termed replication catastrophe, promotes cell lethality (Buisson et al., 2015).

Potency state appears to impact how cellular pathways manage genomic threats. Naïve cells of the blastocyst are less sensitive to irradiation induced DNA breaks than primed epiblast cells (Heyer et al., 2000). In this context, primed epiblast cells exhibit a greater apoptotic sensitivity to due to differential Atm-p53 activity relative to their naïve counterparts (Laurent and Blasi, 2015). However, while both Atm and Atr are active during pluripotency, only *Atr* is essential (Barlow et al., 1996; de Klein et al., 2000). Lethality of *Atr^-/-^* embryos occurs between E4.5 and E8.5, concomitant with the switch from naïve to primed potency at peri-implantation (de Klein et al., 2000). Further, peri-implantation is the specific developmental window during which many replication stress response factors become essential (Kafer and Cesare, 2020). Thus, we hypothesise that pluripotent cell responses to genomic threats may vary across the potency states of early embryogenesis, potentially in alignment with developmental pressures that arise during specific developmental windows.

Here we investigate how pluripotent embryonic cells manage replication stress. Despite their shared pluripotent characteristic, we find that mESCs and mEpiSCs displayed profound functional differences in the replication stress response. While mEpiSCs leverage Atr-dependent responses to tolerate replication stress and maintain cell division, mESCs readily experience Atm activation, replication catastrophe, and apoptosis under the same replication stress conditions. These data indicate mESCs and mEpiSCs respond distinctly to replication challenges, suggestive of developmentally regulated engagement of genome-integrity pathways during early embryogenesis.

## Results

### Replication stress induces distinct cellular responses in naïve and primed pluripotent cells

Using a chimera approach, we created genetically identical mESCs and mEpiSCs from an A2lox ESC line (Iacovino et al., 2011; Mazzoni et al., 2011) that are respectively an *in vitro* representation of the naïve inner cell mass cells (~E3.5) and the primed epiblast (~E7.5) (Osteil et al., 2016) (Fig. 1A). Our cultured mESCs and mEpiSCs exhibited typical *in vitro* characteristics including mESC growth in dense small and round colonies, and mEpiSCs proliferation in flatter and sparser colonies (Fig. 1B). To examine the impact of potency state on the cellular response to genome threats, we subjected mESC and mEpiSC cultures to DNA breaks induced by irradiation (IR), and pharmacologically induced replications stress through treatment with the DNA polymerase inhibitor Aphidicolin (APH) or the ribonucleotide reductase inhibitor Hydroxyurea (HU). Measuring apoptosis in live cells with the fluorescent NucView-488 caspase-3 substrate (Smith et al., 2012) revealed subtle increases in apoptosis rates in irradiated mESC and mEpiSC cultures, with mEpiSCs responding slightly more to IR than the mESCs (Fig. 1C and Fig S1A). Replication stress, however, induced substantially differently apoptotic outcomes between the cell lines. Both APH and HU treatments significantly increased apoptosis in mESCs, but not in mEpiSCs (Fig. 1D, E and Fig S1B-E). In agreement, the apoptosis marker cleaved caspase-3 was elevated in whole cell extracts derived from APH or HU treated cultures of pluripotent mESCs, but not mEpiSCs (Fig 1F).

**Figure 1:**
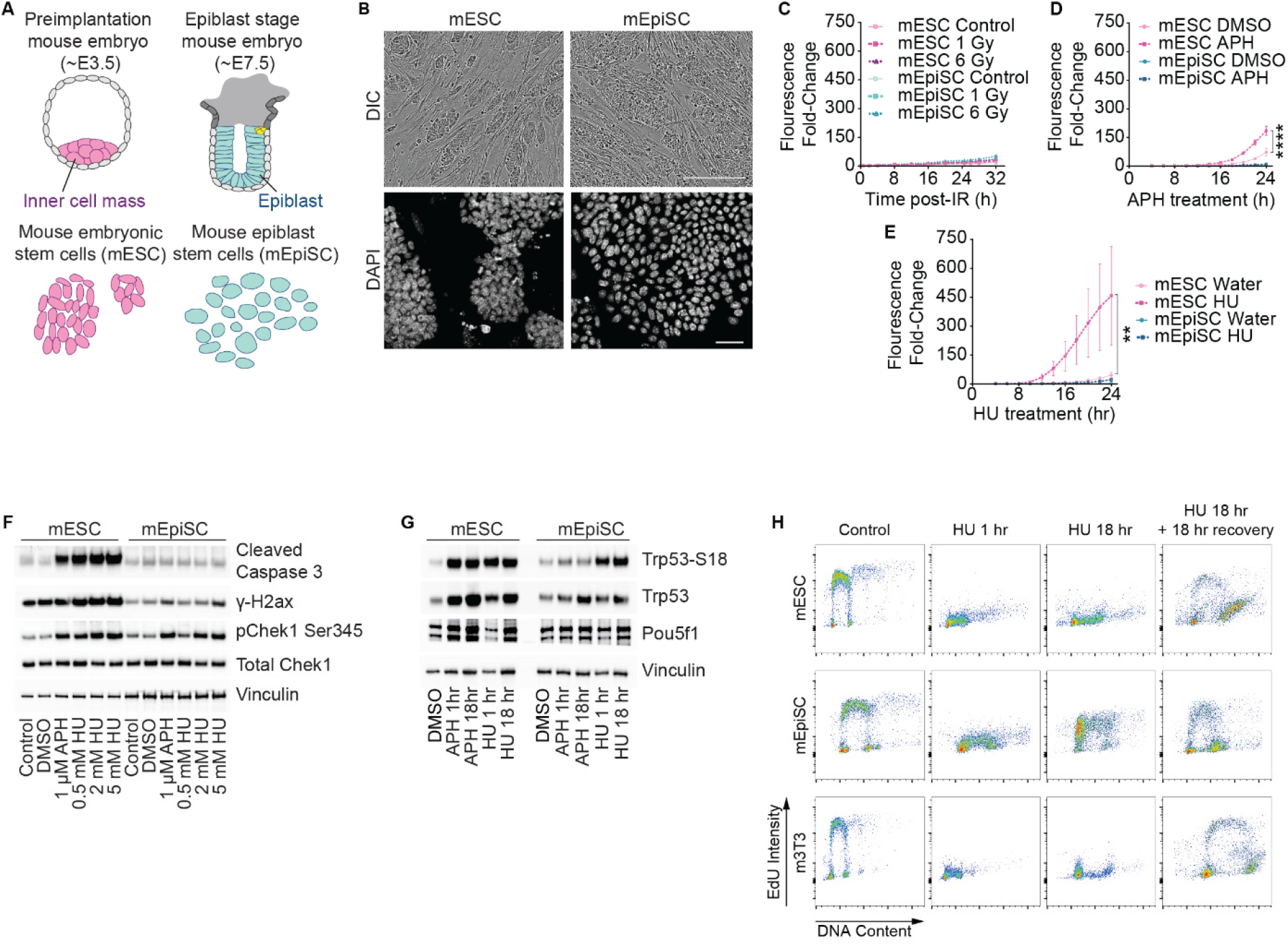
Naïve mESCs but not primed mEpiSCs readily succumb to replication stress. (A) Schematic of sources of murine embryonic (mESC) and epiblast (mEpiSC) stem cells. (B) Differential interference contrast (DIC) and DAPI fluorescent micrographs of stem cells in culture. Scale bar 300 μm. (C-E) Quantification of apoptosis in cultures treated with (C) gamma-irradiation, (D) 1.25 μM Aphidicolin (APH), or (E) 2.5 mM Hydroxyurea (HU). Mean data (± SD) from > 2 independent experiments. Analysis using ANOVA (Bartlett’s test, comparing area under the curve for each condition). Supplementary Figure 1A shows the data from (C) with a different y-axis scale. (F) Western blots of whole cell extracts derived from cultures treated with APH or HU for 18 hours (n > 2 experimental repeats). (G) Western blots on whole cell extracts derived from cultures treated with 2 mM HU for 1 or 18 hours (n > 2 experimental repeats). H) Flow cytometry measuring DNA synthesis (EdU Intensity) and content of cultures treated with 2 mM HU for 1hr, 18 hrs, or 18 hrs followed by an 18-hr recovery period (> 2 experimental replicates).

Following an HU or APH challenge, mESCs exhibited marked increases in phosphorylation of H2ax (γ-H2ax) (Fig 1F and Fig S1F) and Trp53 (p53)-S18 (Fig 1G), and a sustained HU-induced S-phase arrest (Fig 1H, Fig S1G). In contrast, the extent of γ-H2ax and Trp53-S18 phosphorylation in EpiSCs was slightly elevated with HU or APH but did not approach that observed in identically treated mESC cultures (Fig 1 F, G and Fig S1F). Furthermore, mEpiSCs initially halted in S-phase, but eventually overcame HU treatment and resumed replication (Fig 1H).

The different response between potency states was not the result of a blunted DNA damage response (DDR) in either cell line. Both mEpiSCs and mESCs exhibited HU-induced Chek1 (CHK1) phosphorylation, indicative of Atr kinase activity (Fig 1F). Likewise, both cell lines demonstrated similar IR dose-dependent γ-H2ax staining, and phosphorylation of the downstream Atm effector Chek2 (CHK2) (Fig S1G, H). Distinct potency-associated responses also did not result from differentiation in one or both cell lines. Both cell types remained pluripotent in our analysis window as demonstrated by continued Pou5f1 (Oct3/4) expression following APH or HU treatment (Fig. 1G). We focused on HU-induced replication stress for the remainder of this study as reduced nucleotide availability better mimics physiological replication disturbances occurring through high proliferative pressure (Vesela et al., 2017).

### Pluripotency state influences genomic stability following replication stress

Replication stress can manifest as chromosomal aberrations, mitotic errors, and DNA breaks (Wilhelm et al., 2020). Consistent with the observed differential cellular response to replication stress, chromosome breaks were significantly elevated in mitotic mESCs, but not in mEpiSC, following HU treatment (Fig 2A, B). Similarly, HU treated mESCs, but not mEpiSCs, exhibited a significant increase in mitotic separated sister chromatids (Fig 2A, B). This is consistent with premature uncoordinated loss of sister chromatid cohesion, termed cohesion fatigue (Daum et al., 2011); a phenomenon associated with replication stress-induced lethality (Masamsetti et al., 2019). Neutral comet assays revealed that neither cell type exhibited extensive DSBs following HU treatment, though both potency types were equally affected by IR (Fig S2A, B). However, alkaline comet assays, which assess a greater range of DNA damage (Langie et al., 2015), revealed mESCs accrue damage at the low dose of 0.25 mM HU, whereas mEpiSCs require a higher dose of 2 mM HU to induce similar DNA lesions (Fig 2C, D). Under replication stress conditions, the most likely factor promoting elevated tail moments in alkaline but not neutral comet assays is the production of excessive ssDNA associated with challenged DNA synthesis. Collectively the data demonstrate that naïve mESCs are more susceptible than primed mEpiSCs to the negative molecular and cellular outcomes of replication stress.

**Figure 2:**
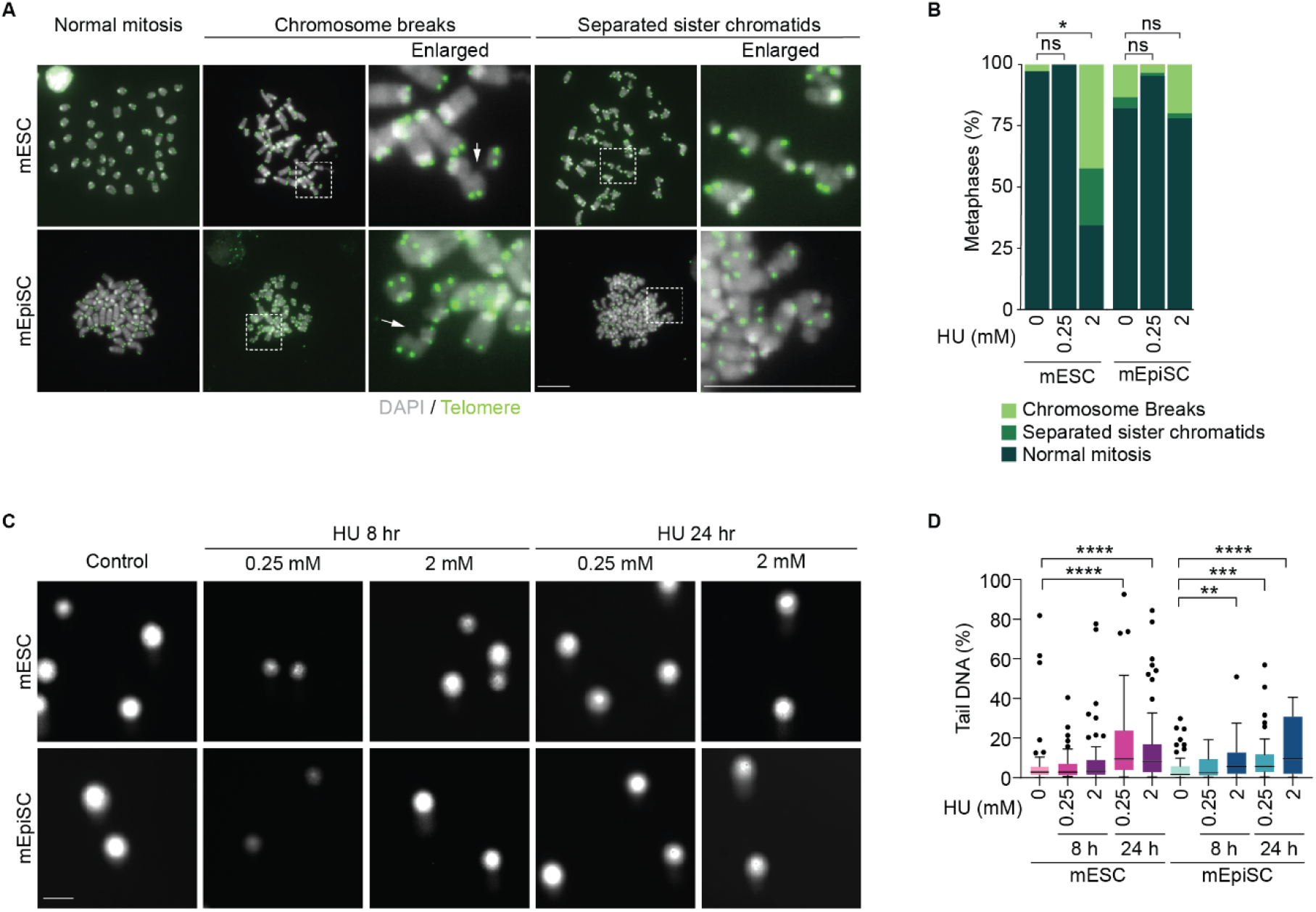
Molecular outcomes to replication stress differ between naïve and primed pluripotent cells. A) Representative images of mitotic chromosome spreads following 24 hours ± 0.25 mM or 2 mM HU. Stained with DAPI (grey) and telomere PNA fluorescent *in situ* hybridization (green), Scale bar = 20 μm. B) Quantification of the experiment in (A) (mean n = 3 experimental replicates quantifying ≥ 15 mitoses spreads per replicate). Raw data was normalized to represent metaphase percentage. P values were calculated using a Wilcoxon test and corrected using the Benjamini – Hochberg method to account for false discovery rate. C) Alkaline comet assay of cultures treated with HU for 8 hrs or 24 hrs, Scale bar 50 μm. D) Quantification of % Tail DNA from (C) in cells treated with HU for 8 or 24 hours (n = 3 experimental replicates of ≥ 30 cells per replicate compiled into a Tukey box plot. Analysis using Kruskal-Wallis and Dunn’s multiple comparisons test. For all panels **** = p < 0.0001, *** = p < 0.001, ** = p< 0.01, * = p < 0.05, ns = not significant.

### Potency state correlates with differential DDR engagement upon replication stress

DDR pathways depend heavily upon the master regulatory kinases Atm and Atr (Blackford and Jackson, 2017). We profiled differences in kinase activity in pluripotent cells under replication stress conditions by performing an unbiased phospho-proteomic analysis of HU-treated mESC and mEpiSC cultures (Fig. 3A). 2,192 significant phosphorylation sites were hierarchically clustered by normalised expression, revealing five primary row clusters (Fig. 3B, Table S1). Kinase motifs were assigned to each phosphorylation site based on the adjacent amino acid residues. The kinases displaying the lowest false-discovery rate across the dataset using Fisher’s Exact test were: Atm/Atr, Prkdc (DNA-PKcs), Gsk3-Erk1-2, Cdk5-Cdk and Cdc2b (Fig. 3C); kinases respectively belonging to the DDR, stem cell potency, and cell cycle control (Blackford and Jackson, 2017; Pao and Tsai, 2021; Singh et al., 2012). We note that Gsk3-Erk1-2 phosphorylation differences were expected given the maintenance of primed and naïve stem cell states requires culture modulation of FGF, activin, and LIF, which signal through Gsk3-Erk1-2 pathways (Cho et al., 2012).

**Figure 3:**
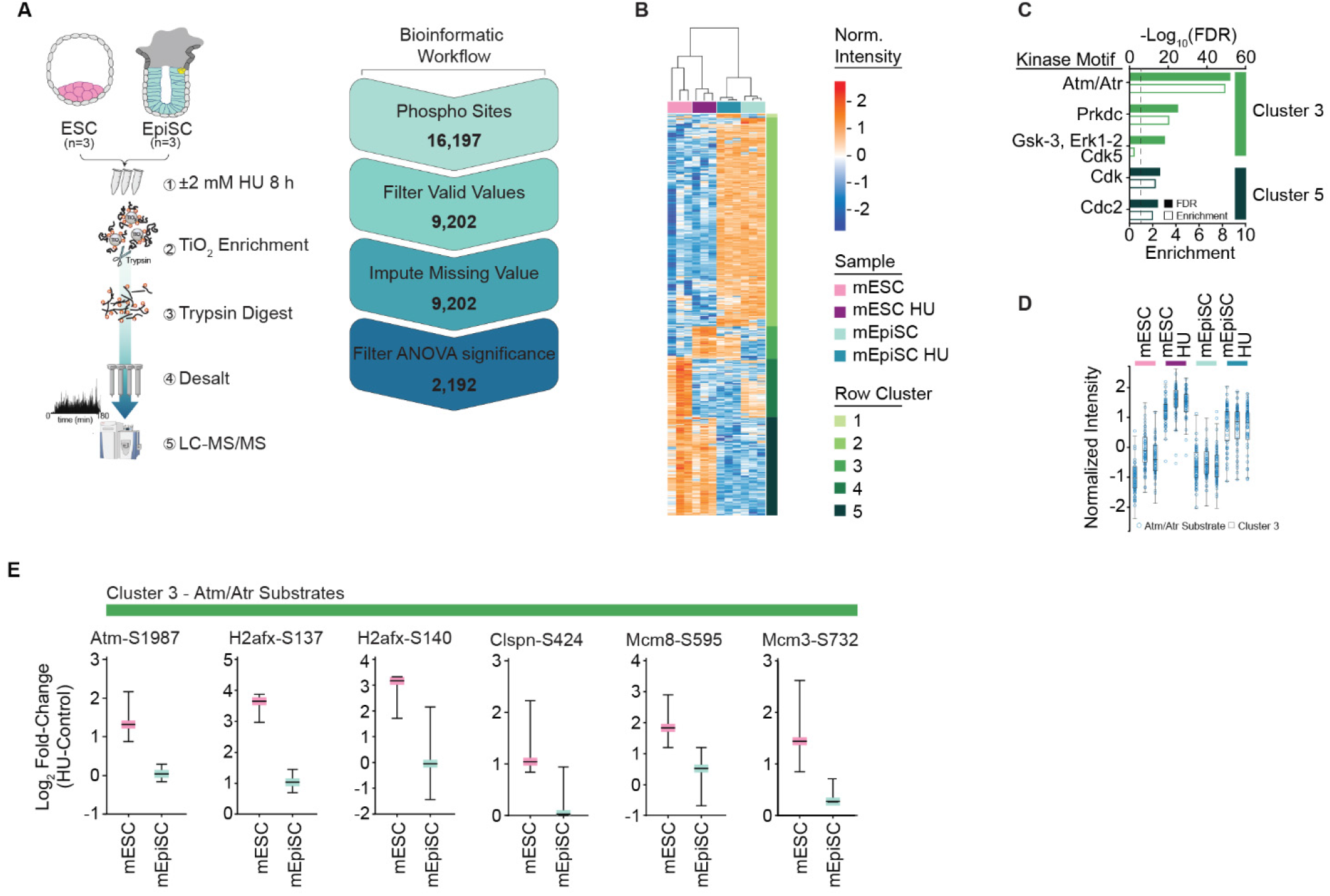
Replication stress induces differential DNA damage response signalling in naïve and primed pluripotent cells. A) Schematic of the mass spectrometry workflow. B) Hierarchically row and column clustered heatmap showing normalized intensity of 2,192 significant phosphopeptides detected by mass-spectrometry in vehicle and HU-treated mESC and mEpiSC cultures (n=3 experimental replicates). C) The top 5 predicted kinase motifs within all row clusters from (B). Rankings determined by −Log_10_FDR, where enrichment values ≥ 1 indicate positive enrichment, analysis using Fisher’s Exact Test. D) Normalised intensity of Atm/Atr substrates from cluster 3, superimposed over all cluster 3 phosphopeptides. E) Log_2_ Fold-Change (HU vs. vehicle) values for a selection of Atm/Atr substrates from cluster 3.

Phosphorylation sites in cluster 3 demonstrated strong positive enrichment of Atm/Atr motifs and were generally highest in the HU-treated mESC samples (Fig. 3C-D). Atm and Atr kinases display overlapping kinase motifs (pS/T-Q) making their substrates difficult to segment in unbiased datasets (Mu et al., 2007). However, a selection of Atm/Atr substrates from cluster 3 were more responsive to HU in mESCs based on their higher Log2 fold-change (Fig. 3E). This included increased phosphorylation of Atm autoactivation site (Atm-S1987) specifically in mESCs cultures. (Fig. 3E). These data were surprising because Atr activity during somatic replications stress (Toledo et al., 2013) suppresses Atm activation, suggesting a different mechanism may be active during replication stress in mESCs.

### Atr suppresses replication catastrophe in primed but not naïve pluripotent cells

To investigate the interplay between Atm and Atr signalling, we treated mESC and mEpiSC cultures with HU and suppressed Atr activity with the specific inhibitor VE-822 (Charrier et al., 2011). Cultures were treated with 2 mM HU for 8 hours ± VE-822, with CldU supplemented for the final 30 minutes to identify replicating cells. In these conditions, γ-H2ax levels in CldU-positive mESCs were significantly increased with HU but were not further elevated with VE-822 co-treatment (Fig. 4A, B). In mEpiSCs, HU alone did not significantly increase γ-H2ax levels, and it was only when cultures were co-treated with HU and VE-822 that H2ax phosphorylation was significantly elevated (Fig 4A, B).

**Figure 4:**
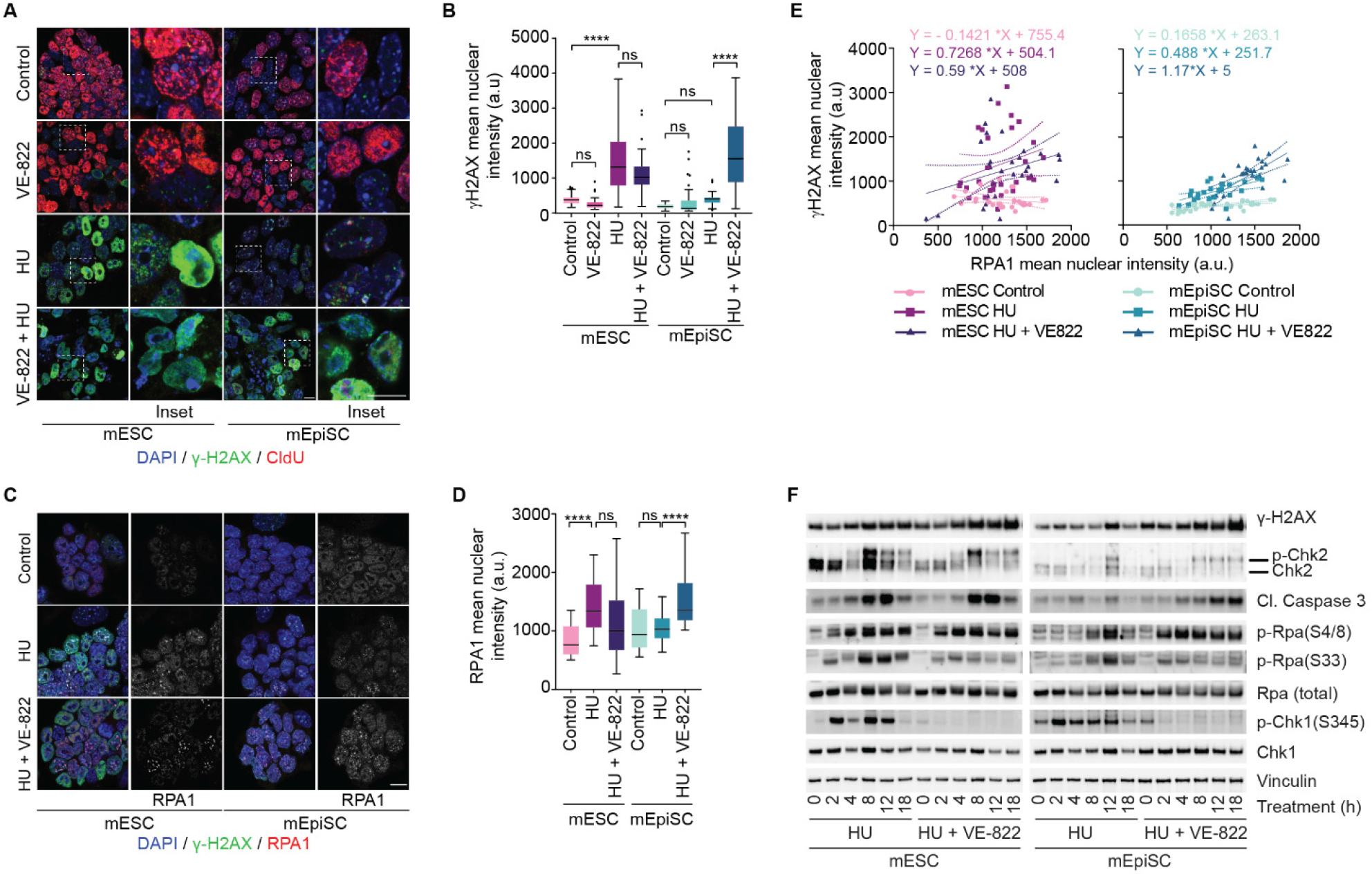
Atr prevents replication catastrophe in primed but not naïve embryonic pluripotent cells. A) Representative micrographs of mESC and mEpiSC cultures treated with, 2 μM VE-822 and 2 mM HU, alone or in combination, for 8 hrs and stained for DNA (DAPI, blue), γH2ax (green), and CldU (red). Scale bar in = 5 μm. B) Quantification of γH2AX mean nuclear intensity in arbitrary units (a.u.) (n = 2 experimental replicates quantifying ≥ 10 nuclei per replicate compiled into a Tukey box plot, analysis performed using ANOVA with Brown-Forsythe post-hoc test, **** = p < 0.001, ns = not significant). C) Representative micrographs of cultures treated with 2 mM HU or HU and 2 μM VE-822 in combination, for 8 hrs, then pre-extracted and stained for DAPI (blue), γH2ax (green) and Rpa1 (red). Scale bar = 5 μm. D) Quantification of Rpa1 mean nuclear intensity (a.u.) from the experiment in (C), n = 2 experimental replicates quantifying from ≥ 10 nuclei per treatment, data presentation and statistics as described in (F). (E) Correlation plots (linear regression) of γH2ax and Rpa1 mean nuclear intensity from mESC and mEpiSC from (C). Linear regression equations for each slope are indicated. (F) Immunoblots of whole-cell extracts derived from cultures treated with 2 mM HU alone or in combination with 2 μM VE-822 for the indicated time (representative example of 2 experimental repeats). For all panels **** = p < 0.0001, ns = not significant.

In somatic and cancer cells, Atr activation prevents replication catastrophe (Toledo et al., 2013). During replication stress, Atr activity prevents generation of excessive ssDNA at stalled replication forks. When Atr is inhibited, replication stress continues to generate ssDNA that bound by Rpa, resulting in the eventual depletion of the cellular Rpa pool. Continued production of ssDNA leads to exposed single-strand segments, DNA breakage, Atm-activation, and a consequential increase in γ-H2ax (Toledo et al., 2013). Chromatin bound Rpa1 increased significantly in mEpiSCs when cells were co-treated with HU and VE-822, but not when treated with HU alone (Fig 4C, D). Conversely, mESCs accumulated chromatin bound Rpa1 when treated solely with HU, and Rpa1 accumulation did not increase further with VE-822 co-treatment (Fig 4C, D). Replication catastrophe is detected by co-immunofluorescence of chromatin bound Rpa1 and γ-H2ax (Toledo et al., 2013). We found that replication catastrophe occurs in mESCs treated with HU alone or HU + VE-822 (Fig 4E). While mEpiSCs accumulated both Rpa1 and γ-H2ax exclusively in HU + VE-822 conditions, and even then, to a lesser degree than observed in mESCs (Fig 4E). The data suggest mESCs are primed to enter replication catastrophe when faced with replication challenges, while mEpiSCs leverage Atr-dependent mechanisms to retain viability during replication stress.

Atr activation is signalled through phosphorylation of Rpa2-S33 and Chek1, whereas Atm phosphorylates Rpa2-S4/8 and Chek2 (Liu et al., 2012). Both mEpiSCs and mESCs display increased Atr activation with HU, and consequential Atr inhibition with VE-822 co-treatment. Replication catastrophe corresponds with Atm-dependent signalling as DNA breaks accumulate (Toledo et al., 2013). In congruence with the above observations, Atm-dependent phosphorylation of Chek2 and Rpa2-S4/8 occurred in mEpiSCs treated with HU + VE-822 but not HU alone. Conversely, Chek2 and Rpa2-S4/8 were readily phosphorylated in mESCs with HU alone and with HU + VE-822 co-treatment (Fig 4F).

Both mESCs and mEpiSCs retained pluripotency in our analysis window during Atr inhibition (Fig S3A), suggesting the observed differences in replication catastrophe were not due to induced differentiation of either cell line. Further, consistent with replication catastrophe induced death, elevated levels of the apoptotic marker cleaved caspase-3 were evident in both HU and HU + VE-822 treated mESCs, but were only elevated in mEpiSCs when cultures were cotreated with HU and VE-822 (Fig 4F, Fig S3A). Both cell types activated and accumulated Trp53 following replication stress (Fig 1G). However, p21, a p53-regulated downstream mediator of cell cycle arrest (Abbas and Dutta, 2009) accumulated more in HU treated mEpiSCs than mESCs (Fig S3A,B). This is consistent with mEpiSCs arresting proliferation in response to replication stress while mESCs instead promoted cell death. In different mESC and mEpiSC cell lines [R1 mESCs (Nagy et al., 1993) and Delmix mEpiSCs] we observed elevated cleaved caspase-3 in both the HU and HU + VE-822 treated R1 cultures, but only in the in the HU + VE-822 treated Delmix cells (Fig S3C). Together the data indicate that under identical replication challenges, primed epiblast cells counter replication stress using Atr-dependent signalling to maintain viability, whereas naïve mESCs readily experience replication catastrophe and induce apoptosis.

## Discussion

Our findings demonstrate different tolerance and management of replication stress between naïve and primed pluripotent cells. When subjected to identical pharmacological replication stress, naïve mESCs displayed more frequent chromosome segregation errors, required a lower HU dose to induce genomic damage, displayed a greater inability to resume DNA synthesis, and presented higher levels of cleaved Caspase-3 than chimera-derived and genetically matched primed mEpiSCs. The Atr/Chek1 and Atm/Chek2 pathways were respectively activated with replication stress and IR in both potency states. However, a distinction between the cell lines was drawn regarding the impact of Atr activity on cell and molecular outcomes. In HU treated mEpiSC cultures, Atr prevented excessive Rpa1 chromatin loading; Rpa, H2ax, and Chek2 phosphorylation; and apoptosis. Conversely, Rpa1 chromatin loading; Rpa, H2ax, and Chek2 phosphorylation; and apoptosis readily occurred in HU treated mESCs regardless of Atr activity. In this regard, the replication stress response of mEpiSCs resembles that of somatic tissues, where Atr was shown previously to prevent replication catastrophe (Toledo et al., 2017; Toledo et al., 2013). In contrast, naïve mESCs differ from other cell types by promptly succumbing to apoptosis in the face of a replication challenge, irrespective of Atr activity.

We do not suggest Atr is absent from the replication stress response in naïve pluripotent cells. Instead, we interpret our findings to indicate naïve and primed pluripotent cells manage replication stress differently. Replication stress is a pervasive threat and cultured mESCs display markers of spontaneous replication defects (Ahuja et al., 2016). This includes H2ax phosphorylation and remodelled replication forks protected by Rad51 (Ahuja et al., 2016). Atr is implicated in somatic replication fork remodelling and protection (Berti et al., 2020), and inhibiting Atr reduces replication rates (Blakemore et al., 2021) and partially supresses spontaneous γ-H2ax (Ahuja et al., 2016) in mESC cultures. Additionally, we observe Atr-dependent Chek1 phosphorylation in HU treated mESCs. What is unique in naïve pluripotent cells is the ease in which cultures activate Atm and slide into replication catastrophe when faced with a replication challenge. Considering this finding, it is not surprising that Atm was recently observed to play a prominent role in the replication stress response of naïve mESC cultures (Blakemore et al., 2021). In agreement, *Atr* deletion is lethal after E4.5, during a later developmental window than blastocyst-derived mESCs, and consistent with a less prominent role for Atr during naïve pluripotency (de Klein et al., 2000).

Despite our efforts we did not identify why replication stress is managed differently in mESC and mEpiSC cultures. However, we interpret these data to indicate that embryonic cells tune replication stress response management to address cell fitness, proliferation demands, and/or developmental milestones associated with each potency state. One possibility is that Atr and replication stress responses are dampened in mESCs to rapidly clear damaged cells. Pre-implantation embryos are generally isolated from exogenous threats, and a diminished replication stress response is likely sufficient to cope with spontaneous replication defects early *in utero.* However, should replication become excessively challenged in the naïve state, Atr is quickly overwhelmed, and replication catastrophe removes the cell in question. This strategy could quickly eliminate potentially damaged cells and prioritize cell quality. At peri-implantation, proliferation rates substantially accelerate and factors critical in the replication stress response become essential during this time (Kafer and Cesare, 2020). We found Atr assumes somatic-like function in primed pluripotent cells and suppresses replication catastrophe. The temporal significance of engaging a complete replication stress response during primed pluripotency is not clear. Primed pluripotent cells may prioritize growth arrest and repair of replication stressed cells to cope with increasing proliferative demands that potentially impart greater endogenous replication stress. Additionally, we speculate that as the embryo moves into gastrulation and organogenesis, the development of more complex embryonic structures benefit from supporting cell fidelity to maintain nascent tissue architecture.

How stem cells maintain genome integrity continues to be explored. Recent data show that telomere protective mechanisms in mESCs diverge from somatic tissues (Markiewicz-Potoczny et al., 2021; Ruis et al., 2021). Here we demonstrated differences in the mESC replication stress relative to mEpiSC and somatic tissues. If, and how, other genome maintenance strategies diverge between somatic and pluripotent tissues is a topic for future exploration.

## Acknowledgements

The Australian Cancer Research Foundation (ACRF) supported Telomere Analysis Center, and Biomedical Proteomics Facility, both located at the Children’s Medical Research Institute (CMRI), are thanked for microscopy and mass spectrometry infrastructure. We thank Pablo Galaviz for his assistance with statistical analyses. GK is supported by an Ideas grant from the Australian National Health and Medical Research Council (NHMRC) (2011344). HAP and laboratory are supported by NHMRC Ideas grants (1187606 and 2003250). PT and laboratory are supported by NHMRC Project Grants (632776, 1127976) and a Research Fellowship (1110751), and by an Australian Research Council (ARC) Discovery Project grant (DP160103651). AJC and laboratory are supported by an ARC Future Fellowship (FT210100858) and Discovery Project grant (DP210103885), an NHMRC Ideas grant (1185870), and philanthropy from the Neil and Norma Hill Foundation.

## Author contributions

GRK and AJC conceived of the study.

GRK, AO’C, PT, AJC designed experimentation.

GRK and AO’C were the primary scientists and completed all cell and molecular biology, live and fixed imaging, and biochemical analysis.

SR completed all mass spectrometry and associated analysis.

PO created cell lines and provided feedback on experimental design.

PG assisted with statistical analysis.

CBN assisted with comet assays and analysis under supervision from HAP.

GRK, AO’C, SR, AJC completed experimental analysis and interpretation and created the figures. GRK, AO’C and AJC wrote the manuscript with assistance from and SR with feedback from all authors.

## Inclusion and diversity

This study did not feature human or non-human subjects, nor genomic datasets. Authors on this paper include one or more individuals who self-identify as an underrepresented ethnic minority in science, who self-identify as LGBTQ+, and who received support from programs designed to increase diversity in science.

## STAR Methods

### Key Resources Tables

| **REAGENT or RESOURCE** | **SOURCE** | **IDENTIFIER** |
| --- | --- | --- |
| *Drugs and chemical reagents* |  |  |
| DMSO | Sigma Aldrich | 276855 |
| Aphidicolin | Sigma Aldrich | A0781 |
| Hydroxyurea | Sigma Aldrich | H8627 |
| VE-822 | SelleckChem | S7102 |
| YOYO-1 | ThermoFisher Scientific | Y3601 |
| EdU | ThermoFisher Scientific | A10044 |
| CldU | Sigma Aldrich | C6891 |
| Telomere PNA probe (Alexa488-OO-(CCCTAA)<sub>3</sub>) | Panagene | F1004 |
| 6-carboxyfluorescein-TEG azide | Berry and Associates | FF 6110 |
| sodium L-ascorbate, | Sigma Aldrich | A4034 |
| Copper (II) sulphate | Sigma Aldrich | 451657 |
| Proteinase K | Sigma Aldrich | 03115852001 |
| Methanol | Sigma Aldrich | 322415 |
| RNase A | Qiagen | 1007885 |
| Triton-100X | Sigma Aldrich | X100 |
| Ponceau S | Sigma Aldrich | P7170 |
| Ethanol | Supleco | 1.00983,2511 |
| EDTA | Chem-Supply | EA022 |
| Sodium Hydroxide | Ajax Finechem | 482 |
| Formamide | Sigma Aldrich | 221198 |
| BSA | Sigma Aldrich | A7906 |
| NuPAGE MES SDS Running buffer | ThermoFisher Scientific | NP0002 |
| Tween-20 | Sigma Aldrich | P7949 |
| NucView Caspase-3 Enzyme Substrate 488 | Biotium | 10402 |
| Acetic acid | Ajax Finechem | AJA1 |
| B-mercaptoethanol | Sigma Aldrich | M3148 |
| SDS | Amresco | 0227 |
| Formaldehyde | Merck | 1.04003.2500 |
| Prolong gold antifade mounting media | ThermoFisher Scientific | P36930 |
| Colcemid | ThermoFisher Scientific | 15212012 |
| B-agarase | NEB | M0392L |
| Clarity Western ECL Substrate | Bio-rad | 1705060 |
| larity Max Western ECL Substrate | Bio-rad | 1705062 |
| PBS | Millipore | 2810305 |
| Coomassie Brilliant Blue | Biorad | 161-0406 |
| Sodium Chloride | Sigma Aldrich | S9625 |
| Immersion oil | Zeiss | 444970-9010-000 |
| Ammonium Bicarbonate | Sigma Aldrich | 09830 |
| Trifluoroacetic Acid (TFA) | ThermoFisher Scientific | 85183 |
| Trypsin / LysC | Promega | VA1061 |
| Ammonia Solution (25%) | Merck Millipore | 5.33003 |
| Sodium Deoxycholate (SDC) | Sigma Aldrich | D6750 |
| Tris(2-carboxyethyl)phosphine (TCEP) | Sigma Aldrich | C4706 |
| 2-Chloroacetimide (CAM) | Sigma Aldrich | 22790 |
| Titansphere Phos-TiO Bulk Material, 10 μm | GL Sciences | 5010-21315 |
| Empore C8 SPE Disks | 3M | 66882-U |
| Attract SPE Bio SDB-RPS Disks | Affinisep | Bio-RPS-M-47.20 |
| Optima MeOH (LC-MS) | Fisher Chemicals | A456-1 |
| ReproSil-Pur C18-AQ, 1.9 μm | Dr. Maisch GmbH | r219.aq. |
| Formic acid | Sigma Aldrich | 5.33002 |
| Acetonitrile | Honeywell | LC015 |
| **Antibodies** |  |  |
| Mouse anti-Chk1 | CST | 2360 |
| Rabbit anti-Chk1Ser345ph | CST | 2341 |
| Mouse anti-Chk2 | CST | 2662 |
| Mouse anti-γH2AX | Millipore | 05-636 |
| Rabbit anti-Cleaved caspase 3 | CST | 9661S |
| Mouse anti-Oct4 | Santa Cruz | sc-5279 |
| Rabbit anti-p21 | Abcam | ab188224 |
| Mouse anti-p53 | CST | 2524 |
| Rabbit anti-p53Ser15ph | CST | 9284 |
| Rabbit anti-Rpa | Novus | NB100-332 |
| Rabbit anti-RpaSer4/8ph | Bethyl | A300-245A |
| Rabbit anti-RpaSer33ph | Bethyl | A300-246A |
| Mouse anti-vinculin | Sigma Aldrich | V9131 |
| Mouse anti- β-actin | Sigma Aldrich | A5441 |
| Rat anti-BrdU | Abcam | Ab6326 |
| Polyclonal Goat anti-rabbit-IgG/HRP | Dako | P0448 |
| Polyclonal Goat anti-mouse-IgG/HRP | Dako | P0447 |
| Goat anti-rat Alexa Fluor 594 | ThermoFisher Scientific | A-11007 |
| Donkey anti-mouse Alex Fluor 488 | ThermoFisher Scientific | A-21202 |
| Donkey anti-mouse Alex Fluor 594 | ThermoFisher Scientific | A-21203 |
| Goat anti-rabbit Alexa Fluor 594 | ThermoFisher Scientific | A-11012 |

| *Software and Algorithms* |  |
| --- | --- |
| **Software / algorithm name** | **Source** |
| ZEISS Zen Black and Zen Blue | Zeiss |
| Metafer 4 | Metasystems |
| Image J / Fiji | NIH, USA |
| FlowJo Version V10 | FlowJo |
| Graphpad Prism v9 | GraphPad Software |
| Cell profiler v2.1.1 | Cell profiler |
| Incucyte Zoom | Santorius |
| Image Lab | Bio-Rad |
| MaxQuant (1.6.0.16) | Juergen Cox |
| Adobe Illustrator 2020 | Adobe |

### Resource availability

#### Lead contact

Further information and requests for resources should be directed to the Lead Contact.

#### Materials availability

This study did not generate new unique reagents.

### Method details

#### Cell culture

Mouse cells (immortalised 3T3 fibroblasts, inactivated MEFs (iMEFs, E13), epiblast stem cells (mEpiSC) and embryonic stem cells (mESC) were grown under physiologically normoxic conditions (10% CO_2_, 3% O_2_ humidified environment at 37°C). iMEF and m3T3 cells were maintained in complete DMEM (10% FBS). mESCs were grown in ESC media (DMEM, 10% ES batch tested FBS (heat inactivated), mouse recombinant LIF (10 ng/mL, Millipore ESGRO^®^ ESG1107) with media exchanged daily and cells passaged at ~70% confluency using 0.05% Trypsin. Both mESC and mEpiSC were passaged onto pre-plated iMEFs growing on gelatinised (0.1% gelatin, Sigma Aldrich, G9391) plates. mEpiSCs were grown in EpiSC media containing (KnockOut DMEM (Gibco, 10829018), 20% KnockOut serum replacement (Gibco, 10828010), supplemented with FGF2 (10 μg/mL, R&D, 233-FB/CF) and Activin (20 μg/mL, R&D, 338-AC/CF). mEpiSC media was exchanged daily and cells passaged at ~80% confluency using sequential collagenase (3mg/mL collagenase Type IV, ThermoFisher, 17104-019) and 0.05% trypsin dissociation steps in the presence of ROCKi (1 μg/mL, Stem Cell Technologies, Y-27632) during passage and for 24 hours after splitting. All media was supplemented with 1x GlutaMAX (ThermoFisher, 35050061), 1x MEM NEAA (ThermoFisher, 11140050) and β-mercaptoethanol (0.05 mM, Sigma Aldrich, M3148). iMEFs were generated from PMEF-CFL EMB Millipore EmbryoMax^®^ Primary MEFs expanded and irradiated with 30 Gy. The following compounds were used in cell treatments: dimethyl sulfoxide (DMSO, Sigma-Aldrich, 276855), Aphidicolin (APH, Sigma-Aldrich, A0781), Hydroxyurea (HU, Sigma-Aldrich, H8627), VE-822 (Selleckchem, S7102), or CldU (Sigma-Aldrich, C6891) for indicated times at indicated concentrations. Where necessary, irradiation was performed using a Gammacell 3000 Elan irradiator for the indicated dosage. DIC images were captured using a EVOS cell imaging system (ThermoFisher).

#### Proliferation and apoptosis assays

Cells were seeded into 96-well plates at 5,000 cells / well with a minimum of 2 technical replicates per condition. After 24 hours, cells were treated with the apoptosis indicator NucView Caspase-3 Enzyme Substrate 488 (1:1000, Biotium, 10402) for 30 minutes. Cells were treated as required and placed in an Incucyte live cell imaging system (Sartoris). Cells were imaged every 2 hours for a maximum of 36 hours without interruption. Cell masks were generated using Incucyte Zoom software was used to estimate changes in cell proliferation (recorded as occupied surface area mm^2^) and apoptosis was estimated measuring the intensity of activated NucView substrate (integrated fluorescence intensity calculated using summed pixel intensity in calibrated units to determine the relative fluorescence intensity as units per image). Once defined, cell mask parameters were used unchanged for all analysed data across each experiment.

#### Immunoblotting

Cells were collected via trypsinisation and counted prior to PBS washing and pelleted via centrifugation (1,000 rpm for 5 minutes) prior to snap freezing. Cells were thawed and lysed in 4x Lithium dodecyl sulfate (LDS) at a concentration of 10,000 cells/μL for 10 minutes at RT with intermittent mixing. Extracts were snap frozen again, thawed and proteins denatured at 65 °C for 10 minutes. Protein extracts were resolved on NuPage Novex Bis–Tris 4–12% gels (ThermoFisher, NP0321BOX) and electrophorized at 200V for 35 minutes at RT or 90 minutes at 4°C for greater separation (pChk2) in 1x NuPAGE MES SDS Running buffer (ThermoFisher, NP0002). Proteins were transferred onto nitrocellulose membranes at 100 V for 1 hour using transfer buffer (25 mM Tris-base (Chem-Supply, TA034), 192 mM Glycine [pH 8.3] (Sigma Aldrich, 410225) with 10% methanol (Sigma Aldrich, Chem-Supply, TA034), 322415)). Transfers were confirmed with reversible Ponceau S (Sigma-Aldrich, P7170) then blocked with 5 % skim milk (for non-phosphorylated protein targets) TBS-T (20 mM Tris (Chem-Supply, TA034), 150 mM NaCl (Sigma Aldrich, S9625), 0.1% Tween 20 [pH 7.6] (Sigma Aldrich, P7949)) or 5% BSA-TBS-T (for phosphorylated protein targets) for 1 hour and probed with primary antibody overnight at 4 °C with gentle agitation. Blots were then washed 5 × 5 minutes in 1x TBS-T, probed with secondary antibodies, washed 5 × 5 minutes with TBS-T and rinsed with dH2O before adding standard (Bio-rad, 1705062) or maximum ECL (Bio-rad, 1705062) for 2 minutes prior to digital visualisation on a BIO-RAD ChemiDoc using Image Lab software. See Supplemental Table 2 for primary and secondary antibody details.

#### Flow Cytometry

Cells were treated with 100 μM EdU (ThermoFisher, A10044) for 30 minutes prior to harvesting by trypsinisation. Cells were centrifuged at 2000 RPM for 5 minutes, washed with PBS and centrifuged again prior to fixation with 70 % ethanol. Pellets were washed with PBS, then blocked with 1 % BSA-PBS. Incorporated EdU was labelled with Click chemistry [(10 μM 6-carboxyfluorescein-TEG azide (Berry and Associates, FF 6110), 10 mM sodium L-ascorbate (Sigma Aldrich, A4034), 2mM Copper (II) sulphate (Sigma Aldrich, 451657)] for 30 minutes in the dark. Cells were washed with 0.5% PBST containing 1 % BSA, RNase (Qiagen, 1007885) treated, and DNA was stained with DAPI (0.1 μg/mL, Sigma Aldrich, 10236276001). Data was acquired on the BD FACSCANTO II and analysed using FlowJo software v10.

#### Comet assay

For both neutral and alkaline comet assays, cells were harvested using trypsin, counted, and diluted to 10,000 cells/mL in low melting agarose (LMA, Trevigen, 4250-050-K) at 37°C at a ratio of 1:10 v/v and added to CometSlide™ (Trevigen, 4250-050-K). Cells were embedded in the LMA and incubated in lysis solution (Trevigen, 4250-050-K) overnight at 4°C. For the neutral comet assay, CometSlides with embedded cells were immersed in neutral electrophoresis buffer for 30 minutes prior to electrophoresis at 1 V/cm^2^ for 45 minutes at 4°C. For alkaline comet assays, CometSlides with embedded cells were immersed in alkaline unwinding solution (200 mM NaOH (Ajax Finechem, 482), 1 mM EDTA (Chem-Supply, EA022)) for 20 minutes at room temperature. Electrophoresis was conducted in alkaline unwinding solution at 1 V/cm for 30 minutes. All slides were washed briefly with dH_2_O, immersed in 70% ethanol for 5 minutes and dried at 37°C. DNA was stained with 1 μM YOYO-1 DNA stain (ThermoFisher, Y3601). Images were captured using Axio Imager.Z2 microscope using a Plan-Apochromat 20x/0.8 M27 air objective, HXP 120 V light source, Axiocam 506 imaging device and appropriate filter cube. Images were processed using ZeissZen Blue software. For analysis, a CellProfiler™ pipeline was developed for semi-automated quantification. DNA was selected as the input image with nuclei as the primary object to be identified. Object diameter range was set to 70 – 1000 pixels to accommodate a range of nuclei sizes. To determine the grading threshold, a global strategy was employed (Otsu method, two class, weighted variance, automatic smoothing, threshold correction factor of 0.7) and lower and upper threshold bounds were set as 0.0 – 1.0 respectively. The pipeline allowed image hole filling in identified objects after thresholding and de-clumping. DNA of each object was then manually edited to separate the DNA in the nucleus and the comet tail. The intensity (integrated density) of the DNA in each segment was measured and exported to excel. Percentage of tail DNA was calculated in excel and data was graphed using GraphPad Prism (v9.0)

#### Chromosome spreads

Cells were treated with 0.2 μg/mL of colcemid (ThermoFisher, 15212012) for 2 hours prior to harvesting. Cells were trypsinised and the reaction was quenched with media containing serum. Cells were incubated at 3:1 dH_2_O: DMEM for 10 minutes at 37 °C. Cells were centrifuged at 1250 RPM for 5 minutes, then fixed with ice cold 3:1 methanol (Sigma Aldrich, 322415):acetic acid (Ajax Finechem, AJA1) overnight at 4 °C. Cells were dropped onto humidified slides which had been washed with 100 % methanol for 1 hour prior and airdried overnight. Slides were rehydrated in PBS for 5 minutes, fixed with 4 % formaldehyde (Merck, 1.04003.2500) /PBS for 5 minutes and washed 3 × 2 minutes with PBS. Slides were treated with 0.25 mg/mL RNaseA (Qiagen, 1007885) / PBS for 15 minutes at 37 °C, fixed with 4 % formaldehyde/PBS for 2 minutes, washed with PBS 3 × 2 minutes, dehydrated in ethanol (Supelco, 1.00.983.2511), and allowed to air dry. A telomere PNA probe (0.3 μg/mL, Alexa488-OO-(CCCTAA)3, Panagene, F1004) was added to the slides, followed by denaturation for 10 minutes at 70°C then hybridised overnight at 37 °C. Slides were washed in PNA wash A (70 % formamide (Sigma Aldrich, 221198), 10 mM Tris pH 7.5) for 2 × 10 minutes followed by PNA wash B (50 mM Tris pH 7.5, 150 mM NaCl, 0.8 % Tween20) 3 × 5 minutes. DNA was stained with DAPI (0.1 μg/mL, Sigma Aldrich, 10236276001) and slides were mounted with DABCO [(2.3% 1,4 Diazabicyclo (2.2.2) octane (Sigma Aldrich, D27802, 90 % glycerol (Sigma Aldrich, G5516), 50 mM Tris pH 8.0 (Amresco, 92161)] anti-fade. Images were captured on a Zeiss AxioImager Z.2 with a 63×, 1.4 NA oil objective and immersion Oil 518F (Carl Zeiss, Wetzlar), appropriate filter cubes and a CoolCube 1m camera (Metasystems). Automated metaphase finding and image acquisition for these experiments were done using Metafer4 v3.12.8, MetaSystems imaging platform.

#### Immunofluorescence microscopy and analysis

All samples were grown on gelatinised (0.1% gelatin) cover glass (thickness No. 1.5H (0.170 mm ± 0.005 mm, Australian Scientific, 0117530) and fixed with 4 % PFA for 10 minutes at 4 °C. Samples processed for replication catastrophe experiments were pre-treated with ice cold 0.2% PBS-TritonX for 1 minute prior to fixation. In experiments assaying CldU thymidine analogue incorporation, cells were pulsed with 50 μM CldU (Sigma-Aldrich, C6891) as indicated prior to fixation. Samples were stored in PBST (0.1 % Tween-20) prior to labelling for no more than 5 days. For labelling, samples were washed 3 x 5 minutes in PBST, permeabilised 1 × 5 minutes in PBS-Triton 100X (0.5 %, Sigma Aldrich, X100), then washed 3 × 5 minutes in PBST. Samples were blocked for 2 hours in 2 % BSA-PBST, then incubated with primary antibody diluted in block solution overnight at 4 °C (see Supplementary Table 2). Samples were washed 5 x 3 minutes in PBST and incubated in the dark with secondary antibody diluted in block solution for 1.5 hours at RT. Samples were washed 5 × 3 minutes in PBST. DAPI (1 μg/ml, Sigma Aldrich, 10236276001) was included in the final wash, before brief rinsing in dH2O and dehydration with sequential 3-minute ethanol washes (70 %, 90 % and 100 %) prior to mounting onto microscope glass (Bio-strategy, EPBRSF41296SP) with Prolong Gold (ThermoFisher, P36930)). Cells were imaged on a Zeiss Airyscan LSM 880 AxioObserver confocal fluorescent microscope fitted with an Airyscan detector using either a 40x or 63x 1.4 NA M27 oil-immersion objective and immersion liquid (Zeiss, 444970-9010-000). Representative images were acquired from a minimum of 5 areas across the coverslip using identical imaging parameters (percent excitation laser power with 1×1 binning for all laser conditions and appropriate filter sets) for like staining in all experiments. For each image, the top and bottom z-plane limits were identified, and z-stack imaging performed to capture the centre of the nucleus. Central z-stack images were used for analysis. For image analysis, immunofluorescence intensity was measured using ImageJ/FIJI (Schindelin et al., 2012). Briefly, outlines around nuclei were created in the DAPI channel, and intensity measured (mean grey value) in channels which pertained to staining of the protein of interest. Randomly selected cell-free areas were measured in each sample image to estimate background fluorescent readings, which were subtracted from the final intensity value for each nucleus. For this analysis, only interphase cells were included. Cells which lacked a discrete nuclei boundary which could not be easily discerned were not included in the analysis.

#### Phosphopeptide Purification

Cells were harvested via trypsinisation methods described above, washed twice with D-PBS and snap frozen as dry pellets prior to phosphopeptide purification as per the EasyPhos workflow (Humphrey et al., 2018). Briefly, pellets were lysed in SDC buffer (containing 10 mM TCEP and 40 mM CAM) and heated for 5 min at 95 °C. Lysates were sonicated at 4 °C in 2 × 10-minute cycles in a bioruptor Plus diagenode at max output. For each sample, 1.25 mg of lysate was removed and digested overnight at 37 °C with 12.5 μg of Trypsin/LysC. Digested phospho-peptides were enriched on TiO2 beads) at 12:1 (bead wt: protein wt). Enriched phosphopeptides were eluted from the TiO2 beads and desalted on house-made SDB-RPS stage tips, dried and then reconstituted in 5 μL of MS buffer.

#### Liquid Chromatography Tandem Mass Spectrometry and Bioinformatics

Samples were loaded on a ~45 cm x 75 μm (ID) fused silica column, packed in-house with 1.9 μm ReproSil AQ C18 particles on a UltiMate3000 UPLC (Dionex) inside a 50 C column oven (Sonation) attached to an ESI-nano-spray source (ThermoFisher). Peptide fractions were separated over a 195 minute gradient consisting of a binary buffer system Buffer A (0.1% formic acid) and Buffer B (0.1% formic acid 90% acetonitrile at a flow rate of 300 nL/min. Elution occurred with a 20 minute loading in 5% buffer B, 150 minute gradient from 5-30% buffer B, final 5 minute elution 30-60% buffer B, and 5 minute column wash in 95% buffer B. Peptides were analysed on a Q-Exactive Plus (ThermoFisher) operating in positive ion DDA mode, with one full scan (300-1650 m/Z, R=35,000 at 200 m/Z) with 3e6 AGC target, and 20 ms IT. The top 10 peptides peaks were submitted for HCD fragmentation (27% NCE) and MS2 (R=35,000 at 200 m/Z) with 1e5 AGC target and 120 ms IT. Centroided Thermo Raw files were analysed in MaxQuant (1.6.0.16) using standard settings and LFQ quantification and searched against the M. musculus Uniprot database (11/2019 release). The phosphopeptide dataset was processed in Perseus (version 16.10.43) (Tyanova et al., 2016). Briefly, the dataset was filtered for common contaminants, reverse identifications and peptides with < 3 valid intensity values from at least one sample group using Perseus (version 16.10.43). Missing intensity values from the remaining peptides were imputed, and significant phosphorylations were identified using ANOVA with permutation-based FDR for multiple corrections. Linear kinase motifs and functional annotations were applied using Perseus’s built-in packages and the PhosphoSite Plus database (Table S1).

#### Statistical analysis and software

Statistical analysis for chromosome spreads and flow cytometry was performed using custom R scripts (please see Data and code availability). We used R version 4.1.0., with the following packages ggplot2, multcomp and tidyverse. All other statistical analysis was performed using GraphPad Prism (v9.0). Figure legends describe error bars, statistical methods, and replicate details for all experiments. No statistical method was used to predetermine sample size. All experiments were repeated at least twice. Figures were prepared using Adobe Illustrator.

## Data and code availability

The mass spectrometry proteomics data have been deposited to the ProteomeXchange Consortium via the PRIDE (Perez-Riverol et al., 2022) partner repository with the dataset identifier PXD032103. R scripts and chromosome spread data are located on GitHub: https://github.com/ChildrensMedicalResearchInstitute/Kafer_OConnor_et_al_2022.git

## Supplementary Information

**Supplementary Figure 1:**
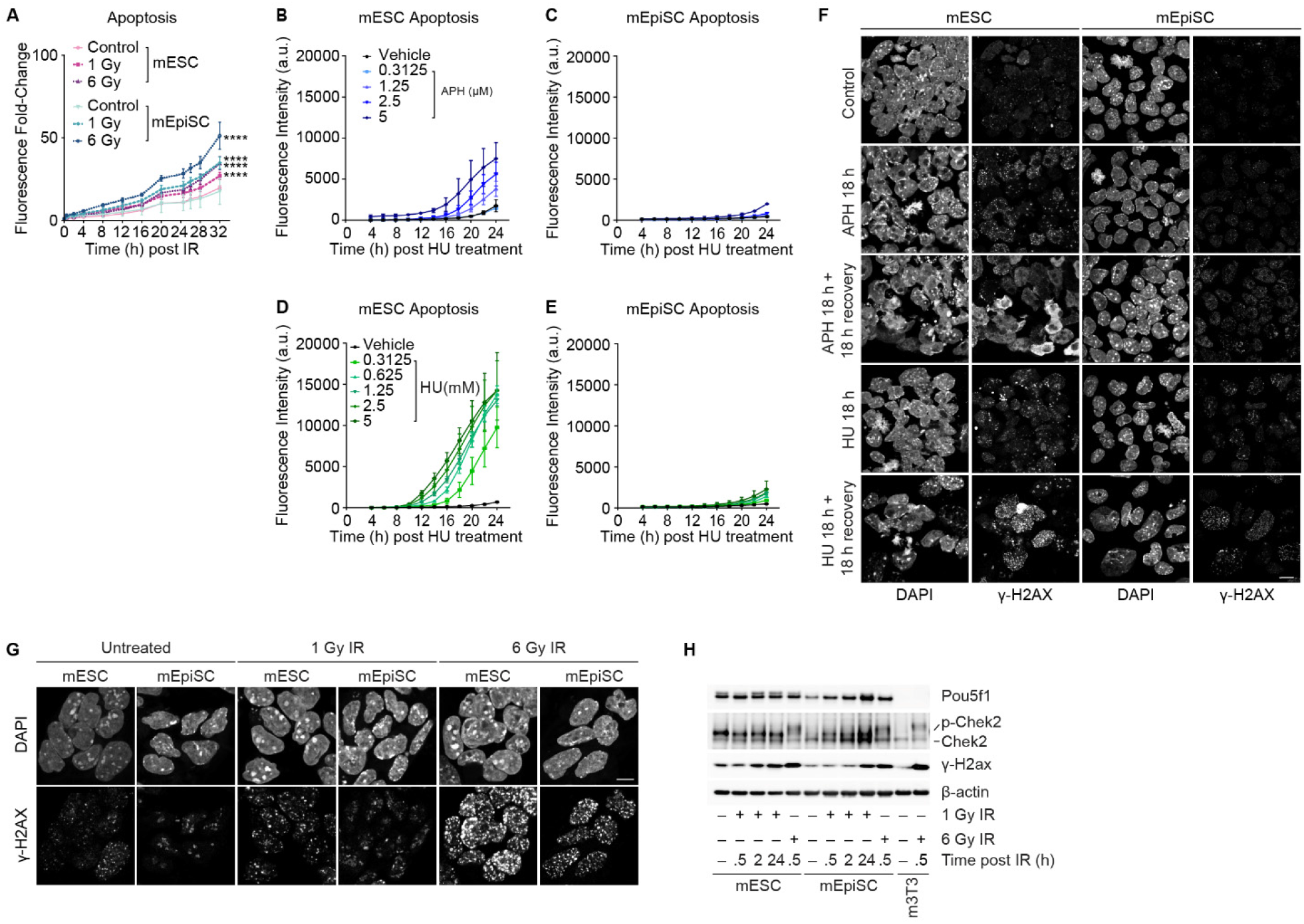
Pluripotent cells of different potency states respond similarly to irradiation, but differently to replication stress. A) Quantification of apoptosis in cultures treated with gamma-irradiation. These are the same data as Figure 1C plotted on a different Y-axis. Mean (± SD) from > 2 independent experiments. ANOVA (Bartlett’s test, comparing area under the curve for each condition). B-E) Quantification of Apoptosis in cultures treated with (B, C) increasing doses of Aphidicolin (APH), or (D, E) increasing does of Hydroxyrera (HU). Mean data (± SD) from > 2 independent experiments. F) Fluorescent micrographs of cultures treated with 1 μM APH or 2 mM HU for 18 hrs, with and without 18 hrs of recovery after treatment (representative images of n > 2 experimental replicates). G) Fluorescent micrographs of cultures fixed 30 minutes after treatment with 1 or 6 Gy IR and stained with DAPI and γ-H2ax (representative images of n > 2 experimental replicates). H) Immunoblots of whole cell extracts derived from cells treated with 1 or 6 Gy IR collected at the indicated times post-treatment (representative blots of n > 2 experimental replicates).

**Supplementary Figure 2:**
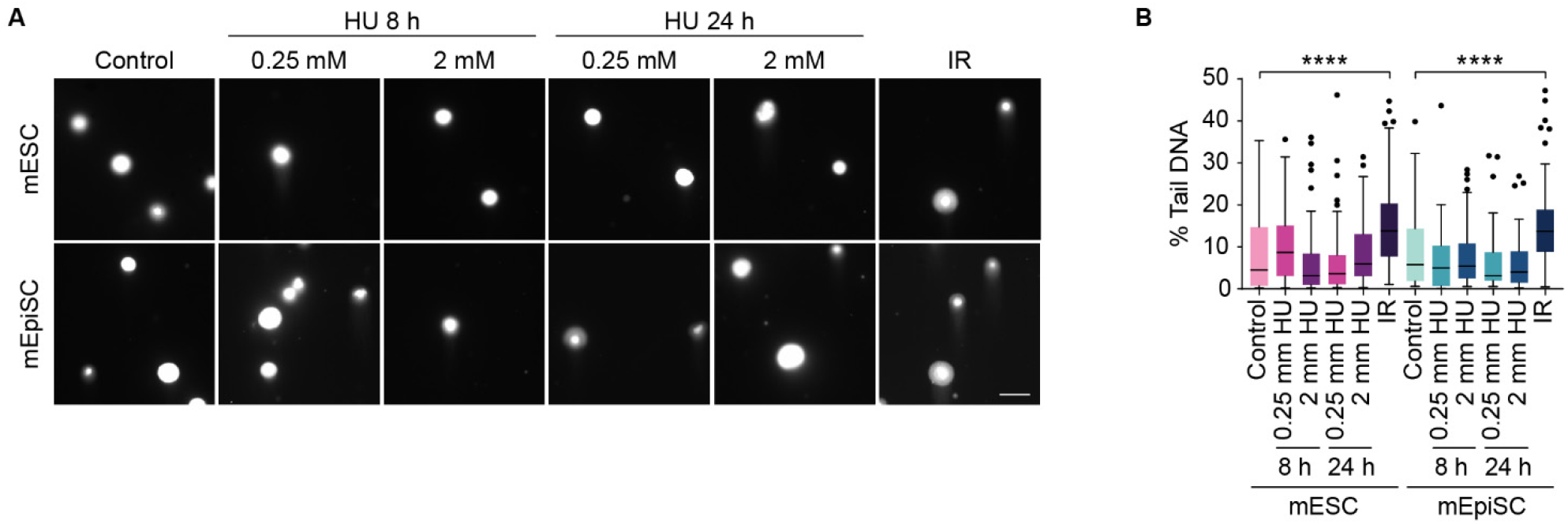
Molecular outcomes to irradiation are similar in naïve and primed embryonic stem cells. A) Representative micrographs of neutral comet assays on cultures treated with 2 mM HU for 8 hrs or 24 hrs, or 1 hr after treatment 10 Gy IR. Scale bar 20 μm. B) Quantification of % Tail DNA from the experiment shown in (A) (n = 3 biological replicates quantifying ≥ 30 cells per replicates compiled into a Tukey box plot, Kruskal-Wallis and Dunn’s multiple comparisons test, **** = p < 0.0001, *** = p < 0.001 and** = p< 0.01).

**Supplementary Figure 3:**
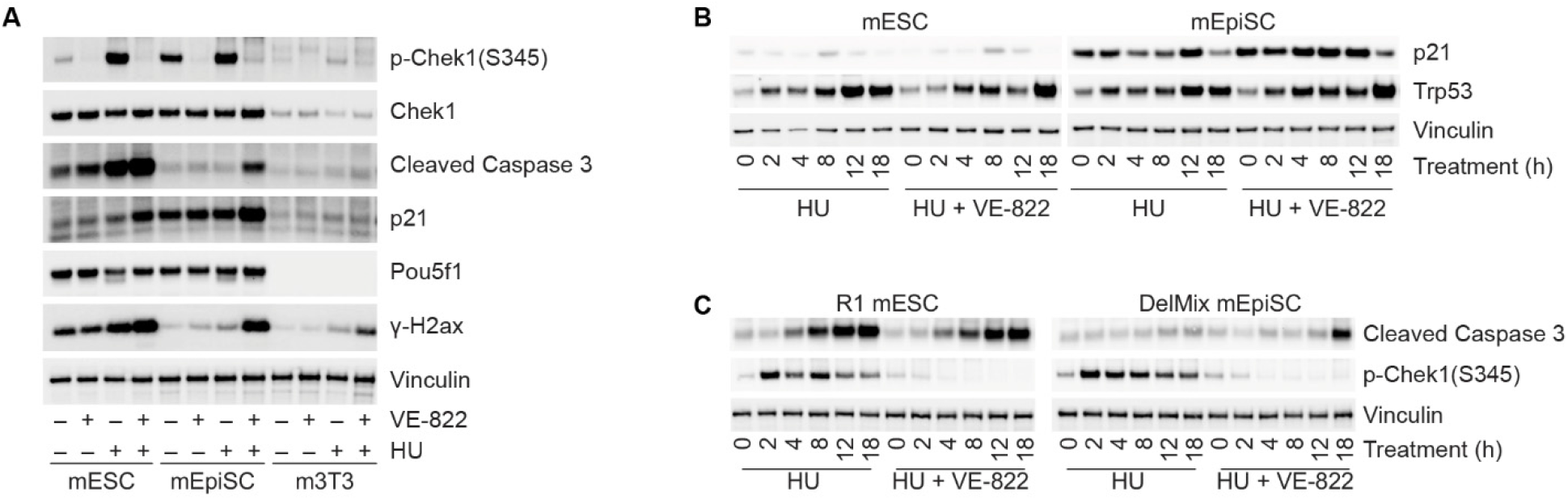
Naïve embryonic stem cells readily undergo replication catastrophe during replication stress. A) Immunoblots of whole cell extracts derived from naïve pluripotent mESC, primed pluripotent mEpiSC, and somatic immortalized m3T3 cultures after 18 hrs treatment with 2 mM HU and 2 μM VE-822, alone or in combination (representative blots from n > 2 experimental replicates). B) Immunoblots of mESC and mEpiSC cultures treated with 2 mM HU alone or in combination with 2 μM VE-822 for the indicated times (representative blots from n > 2 biological replicates). C) Immunoblots of R1 mESC and DelMix mEpiSC cultures after treatment with 2 mM HU alone or in combination with 2 μM VE-822 for the indicated times (representative blots from n > 2 biological replicates).

